# *in-silica* Analysis of SARS-CoV-2 viral strain using Reverse Vaccinology Approach: A Case Study for USA

**DOI:** 10.1101/2020.06.16.154559

**Authors:** Ajay Agarwal

## Abstract

The recent pandemic of COVID19 that has struck the world is yet to be battled by a potential cure. Countless lives have been claimed due to the existing pandemic and the societal normalcy has been damaged permanently. As a result, it becomes crucial for academic researchers in the field of bioinformatics to combat the existing pandemic. The study involved collecting the virulent strain sequence of SARS-nCoV19 for the country USA against human host through publically available bioinformatics databases. Using *in-silica* analysis and reverse vaccinology, two leader proteins were identified to be potential vaccine candidates for development of a multi-epitope drug. The results of this study can provide further researchers better aspects and direction on developing vaccine and immune responses against COVID19. This work also aims at promoting the use of existing bioinformatics tools to faster streamline the pipeline of vaccine development.

**The Situation of COVID19:** A new infection respiratory disease was first observed in the month of December 2019, in Wuhan, situated in the Hubei province, China. Studies have indicated that the reason of this disease was the emergence of a genetically-novel coronavirus closely related to SARS-CoV. This coronavirus, now named as nCoV-19, is the reason behind the spread of this fatal respiratory disease, now named as COVID-19. The initial group of infections is supposedly linked with the Huanan seafood market, most likely due to animal contact. Eventually, human-to-human interaction occurred and resulted in the transmission of the virus to humans. [13].

Since then, nCoV-19 has been rapidly spreading within China and other parts of World. At the time of writing this article (mid-March 2020), COVID-19 has spread across 146 countries. A count of 164,837 cases have been confirmed of being diagnosed with COVID-19, and a total of 6470 deaths have occurred. The cumulative cases have been depicting a rising trend and the numbers are just increasing. WHO has declared COVID-19 to be a “global health emergency”. [14].

**Current Scenario and Objectives:** Currently, research is being conducted on a massive level to understand the immunology and genetic characteristics of the disease. However, no cure or vaccine of nCoV-19 has been developed at the time of writing this article.

Though, nCoV-19 and SARS-CoV are almost genetically similar, the respiratory syndrome caused by both of them, COVID-19 and SARS respectively, are completely different. Studies have indicated that –

“**SARS was more deadly but much less infectious than COVID-19”**.

**-*World Health Organization***

---

The spread of SARS epidemic has provided with any useful insights as during that time the virus epidemic was contained only through general prevention means and treatment of the individual symptoms. As a result, there exists only a limited number of tools available to test the coronavirus for their ability to infect humans. This has acted as a major limitation for predicting the next zoonotic viral outbreak. [13-15].

In response to the current medical crisis, the World Health Organization has activated the R&D Blueprint for the acceleration of the development of diagnostics, therapeutics and vaccines for treatment of this novel coronavirus.

The objective of this study lies in compliance to the guidelines for research activated by WHO. This study aims to utilize a reverse-vaccinology approach in order to identify potential vaccine candidates for COVID19 for the country USA. We shall be deploying open-access bioinformatics tools for our analysis of the same.

## Materials and Methods

### Strain Selection

The virulent strain of SARS-CoV-2 was selected for in-silico analysis. The genome of the viral strain is made available by NCBI - National Center for Biotechnology Information (https://www.ncbi.nlm.nih.gov.in).

The reference identification for SARS-CoV-2 is given by *RefSeq NC_045512*.*2*

### Protein Identification and Retrieval

14534 viral protein sequences of the *SARS-CoV-2* were obtained from the ViPR – Virus Pathogen Database and Analysis. [1]. These protein sequences were identified and downloaded in a tabular format for the *Host – Humans* and the *Country – USA*.

**Table 1.**
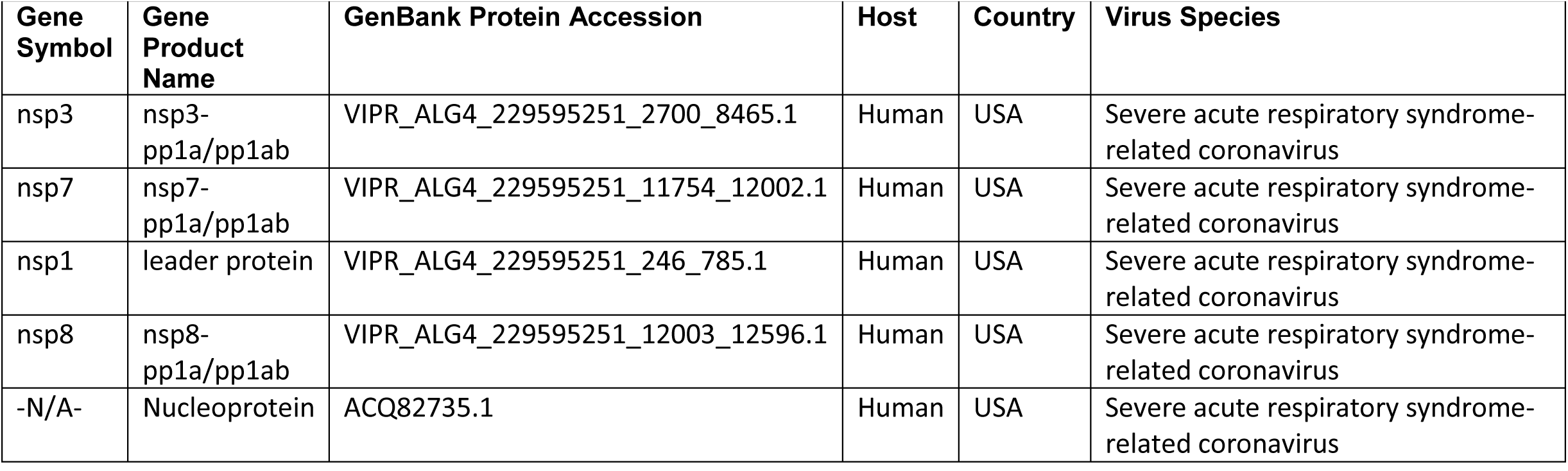
The above table depicts randomly selected four protein sequences obtained out of the 14534 collected under the ViPR databases along with their respective properties. Apart from the properties mentioned in the above table, other parameters obtained from the ViPR databases included the Protein Length, Maturation Status, Collection Data, Entire Protein Sequence Availability, Strain Name etc.

### Physiochemical Property Analysis

The FASTA-file for all the 14535 protein sequences was downloaded and loaded in R using “**SeqinR”** and “**Biostrings”** packages. The different physicochemical properties were analysed using the “**Peptides”** package in the R. [2-4].

### Protein Antigenicity

VaxiJen 2.0 is used to predict the antigenicity of the protein based on the FASTA-files that contain their respective amino acid sequences. The online software predicts the antigenic score for the same. [5-6].

### B-Cell and T-Cell Epitope Prediction

The T-cell and B-cell epitopes of the selected two leader protein sequences were predicted using the IEDB (The Immune Epitope Database). IEDB is a freely available resource funded by NIAID. It is a catalog that contains the experimental data on antibody and T cell epitopes studied in humans, non-human primates, and other animal species in the context of the infectious disease, allergy, autoimmunity, and transplantation. It also assists in hosting tools for predicting and analysing the epitopes. [7].

For MHC Class-I T-cell epitope prediction for ORF1ab leader proteins, NetMHCpan EL 4.0 method was used. This method was applied for the HLA-A*11-01 allele. For MHC Class-II T-cell epitope prediction for ORF1ab leader proteins, Sturniolo prediction method was used. This method was applied for the HLA DRB1*04-01 allele. For B-cell lymphocyte epitope prediction, Bepipered Prediction Method was used. [8].

Ten epitopes were selected on random for analysis of their antigenicity, allergenicity, topology and conservancy.

### Allergenicity, Topology and Toxicity Prediction of the predicted Epitopes

AllerTOP v2 (https://www.ddg-pharmfac.net/AllerTOP/) was used for determining the allergenicity of selected epitopes. ToxinPred server (https://webs.iiitd.edu.in/raghava/toxinpred/protein.php) was used to determining the toxicity of the selected epitopes. The prediction of transmembrane helices in proteins was determined using the TMHMM Server v2.0 (http://www.cbs.dtu.dk/services/TMHMM/).

### Prediction of Conservancy of the Epitopes selected

The conservancy of the selected epitopes was analysed using the conservancy analysis tool of the IEDB server. For this analysis, the parameter for the sequence identity threshold was adjusted to ‘>=50’. [7].

### 3D Structure Generation for Epitopes selected

In order to generate the 3D structure of the epitopes choses, the PEP-FOLD3 tool was used. PEP-FOLD (http://bioserv.rpbs.univ-paris-diderot.r/services/PEP-FOLD3/) uses a *de novo* approach to predict the peptide sequences from the amino acid sequences. [9-11]. It utilizes the structural alphabet SA letters to describe the conformations of four consecutive residues.

### Molecular Docking of the Epitopes selected

The pre-docking of the selected epitopes was done using the UCSF Chimera. It was also used to perform pre-docking of the selected alleles HLA-A*11-01 (for MHC Class-I) and HLA DRB1*04-01.

Later, the docking of the peptide-protein was done using HPEPDOCK. HPEPDOCK is a web server of performing blind peptide-protein utilizing hierarchical algorithm. [12]. Instead of performing length stimulations to refine peptide conformations, HPEPDOCK studies the peptide flexibility through an ensemble of peptide conformations produced by the MODPEP program.

## Results

### Selection and Retrieval of Potential Vaccine Candidate information

The *SARS-CoV-2* strain was identified. All the 14534 protein sequences were analysed on the basis of the physicochemical properties to select the top five candidates for further analysis.

For the physicochemical analysis, the number of amino acids, instability index, aliphatic index, and the grand average of hydrophobicity (GRAVY) scores of the all the 14,534 proteins were taken using the *Peptides* package in R. This packages allows the identification, selection and analysis of multiple amino acid sequences in the same FASTA-file. Hence, it was utilized.

**Table 2.** Physicochemical property analysis for SARS-CoV-2 against the top five viral proteins having the lowest GRAVY scores and confirmed stability out of the 14,534 proteins obtained from ViPR database.

| Ref.No. | Instability Index | Is Stable or Not ? | Aliphatic Index | GRAVY | Gene Symbol | Gene Product Name | GenBank Accession | Strain Name |
| --- | --- | --- | --- | --- | --- | --- | --- | --- |
| 10946 | 29.075 | Stable | 86.5 | -0.43333 | ORF1ab | leader protein | MT326049 | SARS-CoV-2/human/USA/WA-UW-1872/2020 |
| 6333 | 23.70167 | Stable | 76.77778 | -0.42444 | ORF1ab | leader protein | MT326102 | SARS-CoV-2/human/USA/UNKNOWN-UW-1818/2020 |
| 13832 | 27.65111 | Stable | 88.11111 | -0.40167 | ORF1ab | leader protein | MT293180 | SARS-CoV-2/human/USAWA-UW415/2020 |
| 8854 | 28.97056 | Stable | 85.94444 | -0.40111 | ORF1ab | leader protein | MT326175 | SARS-CoV-2/human/USA/WA-UW-1616/2020 |
| 9248 | 27.59556 | Stable | 87.55556 | -0.395 | ORF1ab | leader protein | MT326144 | SARS-CoV-2/human/USA/WA-UW-1695/2020 |

The physicochemical study unveiled five potential candidates having the lowest score of GRAVY and instability index less than 40 (hence, displaying stability). These five candidates were individually analysed for their molar extinction coefficients and antigenicity.

### Antigenicity and Extinction Coefficients of the Potential Candidates

The VaxiJen 2.0 tool was used to analyse the antigenicity of the potential five candidates and the molar extinction coefficient was analysed using the ExPASy’s online tool –*ProtParam*.

Out of the five potential candidates, only two were selected for further analysis. This was based on the criteria of having the highest score of predicted antigenicity and highest values of the extinction coefficients.

**Table 3.** The table depicts the extinction coefficient and antigenicity scores of the selected five potential vaccine candidates from ViPR database.

| Ref No | Extinction Coefficient (in M<sup>-1</sup>cm<sup>-1</sup>) | Antigenicity Scores and its Results |  | Threshold |
| --- | --- | --- | --- | --- |
| 10946 | 12950 | 0.4097 | Antigen | 0.4 |
| **6333** | **12950** | **0.4624** | **Antigen** | **0.4** |
| 13832 | 12950 | 0.4045 | Antigen | 0.4 |
| **8854** | **12950** | **0.5166** | **Antigen** | **0.4** |
| 9248 | 12950 | 0.3886 | Non-antigen | 0.4 |

These two were: ORF1ab leader protein **MT326102** and the ORF1ab leader protein **MT326175**. They both had the same molar extinction coefficient of 12950 but, different yet high scores of predicted antigenicity – 0.4624 and 0.5166 respectively.

Both the leader proteins were then used for further analysis.

### T-cell and B-cell Epitope Prediction

The T-cell epitopes of the MHC Class-I for both the leader protein were determined using the NetMHCpan EL 4.0 prediction method of the IEDB server keeping the sequence length at 9. This tool allows the generation of the epitopes and sorts them on the basis of their percentile scores. Randomly, ten potential epitopes were selected randomly for the *antigenicity, allergenicity, toxicity*, and *conservancy tests*.

For MHC class-II, T-cell epitopes (HLA DRB1*04-01 allele) of the proteins were also determined using the IEDB Analysis tools. The *Sturniolo* prediction methods were used for the same. Again, ten potential candidates were chosen based on the same criteria as that of MHC Class-I.

The B-cell epitopes of the proteins were selected using the Bepipered Linear Epitope Prediction method of the IEDB server.

### Topology Identification of the Epitope

The topology of the chosen epitopes was determined using the TMHMM v2.0 server (http://www.cbs.dtu.dk/services/TMHMM/). The potential T-cell epitopes, whose *topology, antigenicity, allergenicity, toxicity* and *conservancy* was analysed, for ORF1ab leader proteins **MT326102** and **MT326175** are depicted in the following tables.

**Table 4.** List of MHC Class-I epitopes for selected leader protein ORF1ab MT326102 along with their respective antigenicity score, allergenicity, topology, conservancy and toxicity measure. The epitope in bold italics represented the chosen epitope for 3D Generation.

| **Epitope** | **start** | **end** | **Antigenicity and its Scores** |  | **Allergenicity** | **Topology** | **Conservancy** | **Toxicity** |
| --- | --- | --- | --- | --- | --- | --- | --- | --- |
| SLVPGFNEK | 3 | 11 | Antigen | 1.4706 | Allergen | outside | 100 | Non-Toxin |
| VAYRKVLLR | 116 | 124 | Non-Antigen | -0.456 | Non-Allergen | inside | 100 | Non-Toxin |
| HVGEIPVAY | 110 | 118 | Antigen | 0.6413 | Allergen | outside | 100 | Non-Toxin |
| LSEARQHLK | 39 | 47 | Antigen | 0.6432 | Allergen | inside | 100 | Non-Toxin |
| VLLRKNGNK | 121 | 129 | Non-Antigen | -0.7264 | Non-Allergen | inside | 100 | Non-Toxin |
| VGEIPVAYR | 111 | 119 | Antigen | 0.8204 | Allergen | inside | 100 | Non-Toxin |
| GEIPVAYRK | 112 | 120 | Antigen | 0.9683 | Allergen | inside | 100 | Non-Toxin |
| RTAPHGHVM | 77 | 85 | Non-Antigen | 0.2594 | Allergen | inside | 100 | Non-Toxin |
| AYRKVLLRK | 117 | 125 | Non-Antigen | -1.2982 | Non-Allergen | inside | 100 | Non-Toxin |
| **RSDARTAPH** | **73** | **81** | **Antigen** | **0.5509** | **Non-Allergen** | **inside** | **100** | **Non-Toxin** |

**Table 5.**
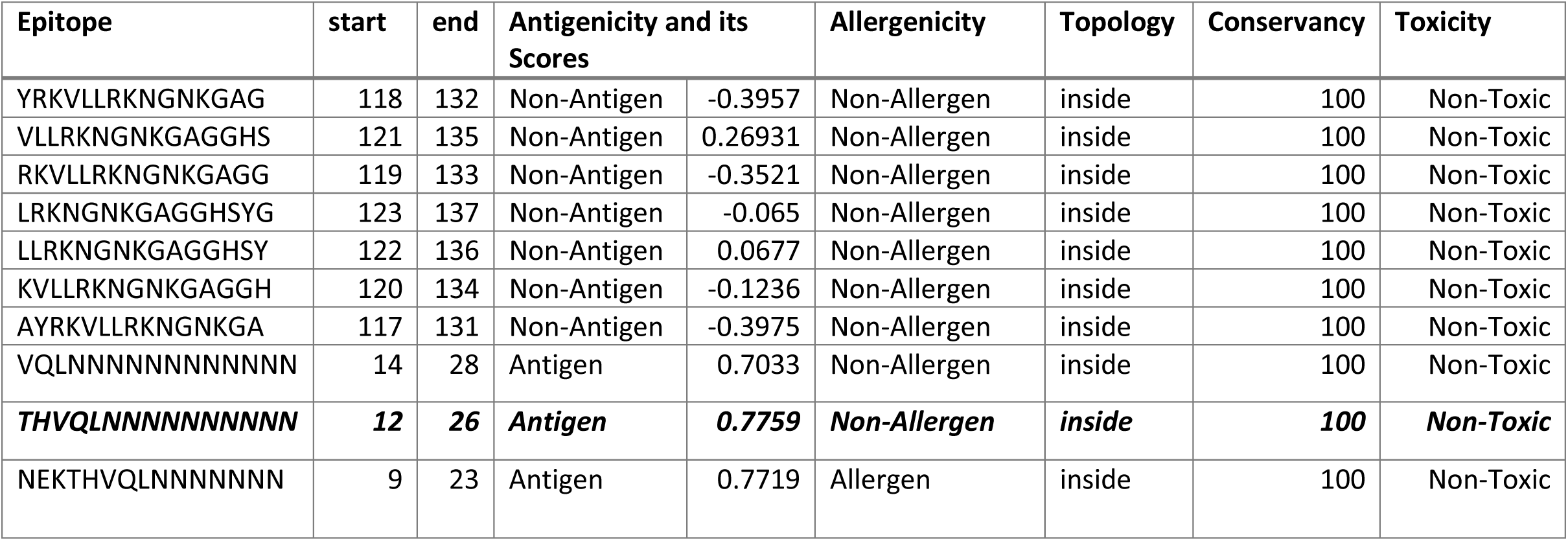
List of MHC Class-II epitopes for selected leader protein ORF1ab MT326102 along with their antigenicity scores, allergenicity, topology, conservancy scores, and their respective toxicities. The epitope in bold italics is selected for further 3D Structure generation.

**Table 6.**
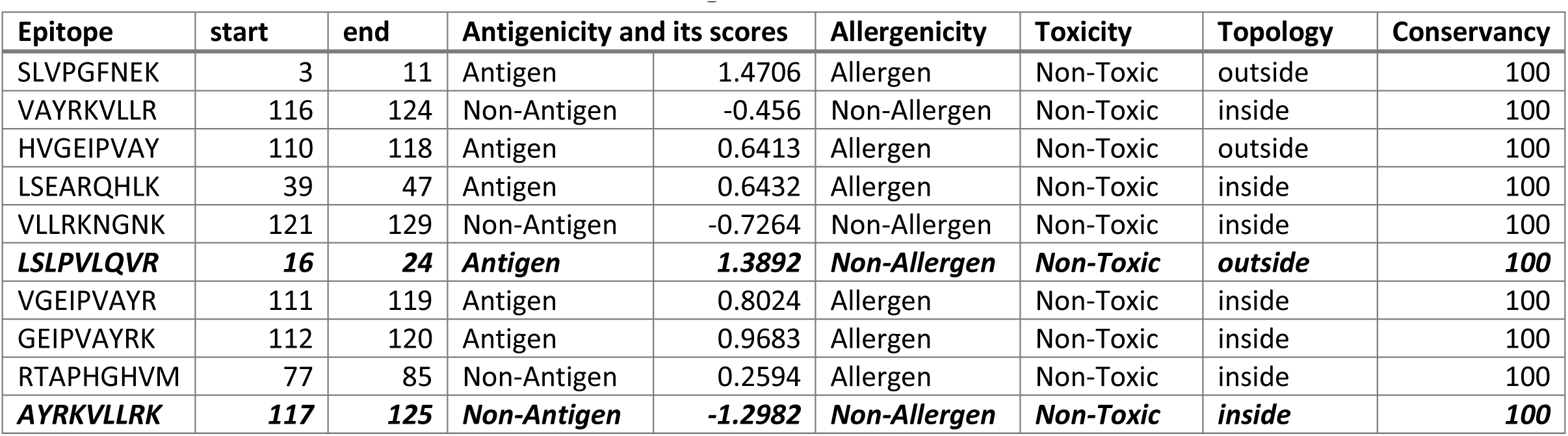
List of MHC Class-I epitopes for selected leader protein ORF1ab MT326175 along with their antigenicity, allergenicity, toxicity, topology and conservancy. The epitope in bold italics is selected for further 3D Structure generation.

**Table 7.**
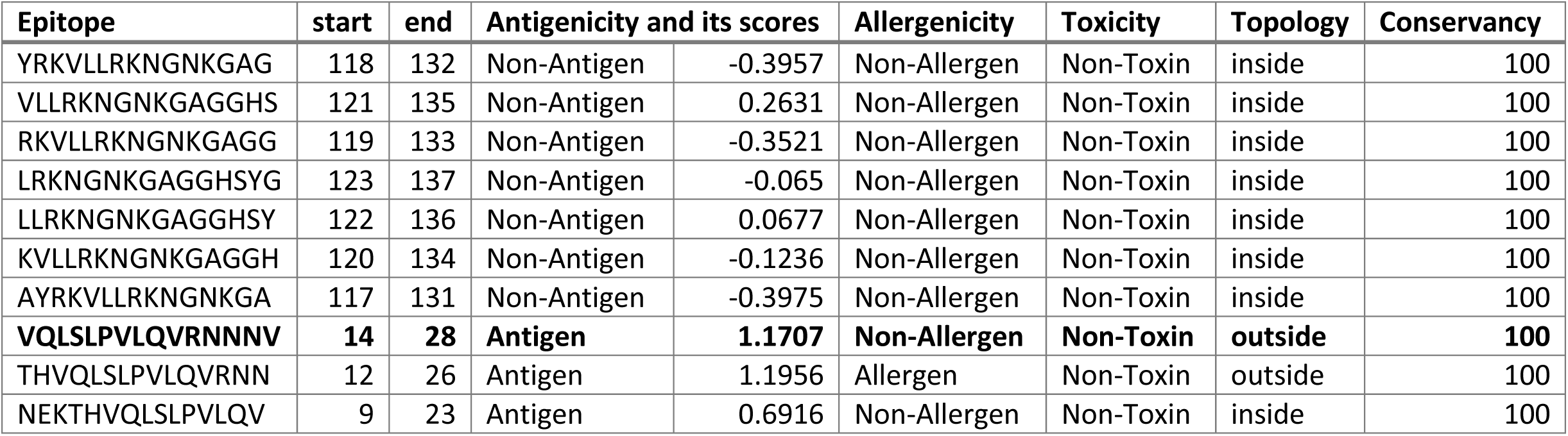
List of MHC Class-II epitopes for selected leader protein ORF1ab MT326175 along with their antigenicity, allergenicity, toxicity, topology and conservancy scores. The epitope in bold italics is used for further 3D Structure generation.

| Epitope | start | end | Antigenicity and its scores |  | Allergenicity | Toxicity | Topology | Conservancy |
| --- | --- | --- | --- | --- | --- | --- | --- | --- |
| YRKVLLRKNGNKGAG | 118 | 132 | Non-Antigen | -0.3957 | Non-Allergen | Non-Toxin | inside | 100 |
| VLLRKNGNKGAGGHS | 121 | 135 | Non-Antigen | 0.2631 | Non-Allergen | Non-Toxin | inside | 100 |
| RKVLLRKNGNKGAGG | 119 | 133 | Non-Antigen | -0.3521 | Non-Allergen | Non-Toxin | inside | 100 |
| LRKNGNKGAGGHSYG | 123 | 137 | Non-Antigen | -0.065 | Non-Allergen | Non-Toxin | inside | 100 |
| LLRKNGNKGAGGHSY | 122 | 136 | Non-Antigen | 0.0677 | Non-Allergen | Non-Toxin | inside | 100 |
| KVLLRKNGNKGAGGH | 120 | 134 | Non-Antigen | -0.1236 | Non-Allergen | Non-Toxin | inside | 100 |
| AYRKVLLRKNGNKGA | 117 | 131 | Non-Antigen | -0.3975 | Non-Allergen | Non-Toxin | inside | 100 |
| **VQLSLPVLQVRNNNV** | **14** | **28** | **Antigen** | **1.1707** | **Non-Allergen** | **Non-Toxin** | **outside** | **100** |
| THVQLSLPVLQVRNN | 12 | 26 | Antigen | 1.1956 | Allergen | Non-Toxin | outside | 100 |
| NEKTHVQLSLPVLQV | 9 | 23 | Antigen | 0.6916 | Non-Allergen | Non-Toxin | inside | 100 |

### Antigenicity, Allergenicity, Toxicity, and Conservancy Analysis of the Epitopes

On the analysis of the antigenicity, allergenicity, toxicity, and conservancy analysis of the T-cell epitopes, it was found that most of them were antigenic, simultaneously being non-allergenic, non-toxic and higher values of conservancy. From the ten selected MHC Class-I and MHC Class-II T-cell epitopes, one from each category were selected from both the leader proteins. The criteria for being selected was having the higher antigenic scores, non-allergenicity, non-toxicity, and conservancy value above 90%.

The selected epitopes were: **RSDARTAPH, VQLNNNNNN, LSLPVLQVR**, and **VQLSLPVLQ**.

The B-cell epitopes of the ORF1ab leader proteins are displayed in the following two figures.

**Figure 1.**
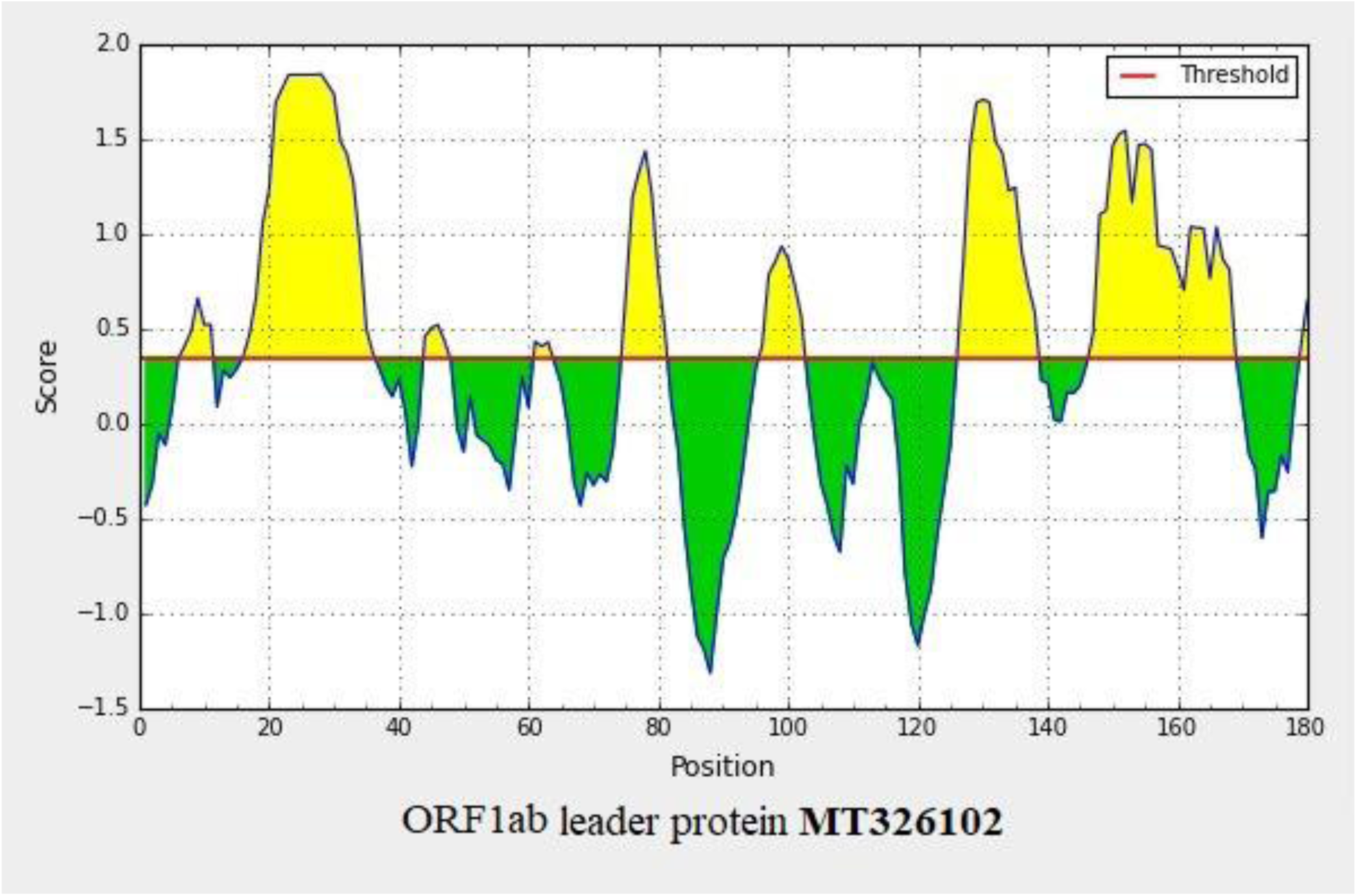
B-cell epitope prediction of the ORF1ab leader protein MT326102 (above). B-cell epitope prediction of the ORF1ab leader protein MT326175 (below).

**Figure 2.**
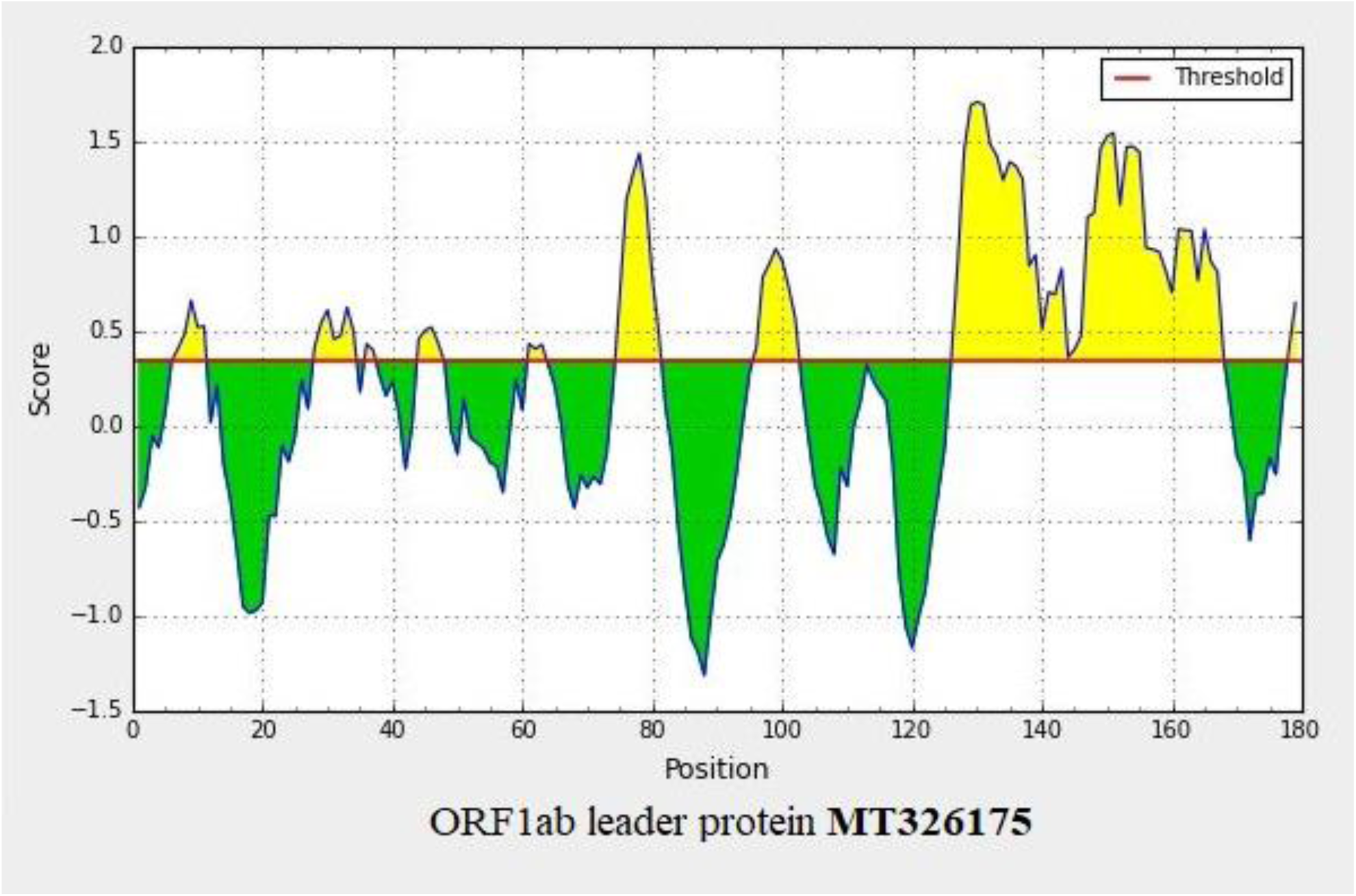
B-cell epitope prediction of the ORF1ab leader protein MT326102 (above). B-cell epitope prediction of the ORF1ab leader protein MT326175 (below).

### Generation of the 3D Structures of Epitopes

The following figures depict the PEP-FOLD3 generated 3D structures of the selected T-cell epitopes of MHC Class-I and Class-II: **RSDARTAPH, VQLNNNNNN, LSLPVLQVR**, and **VQLSLPVLQ**.

**Figure 3.**
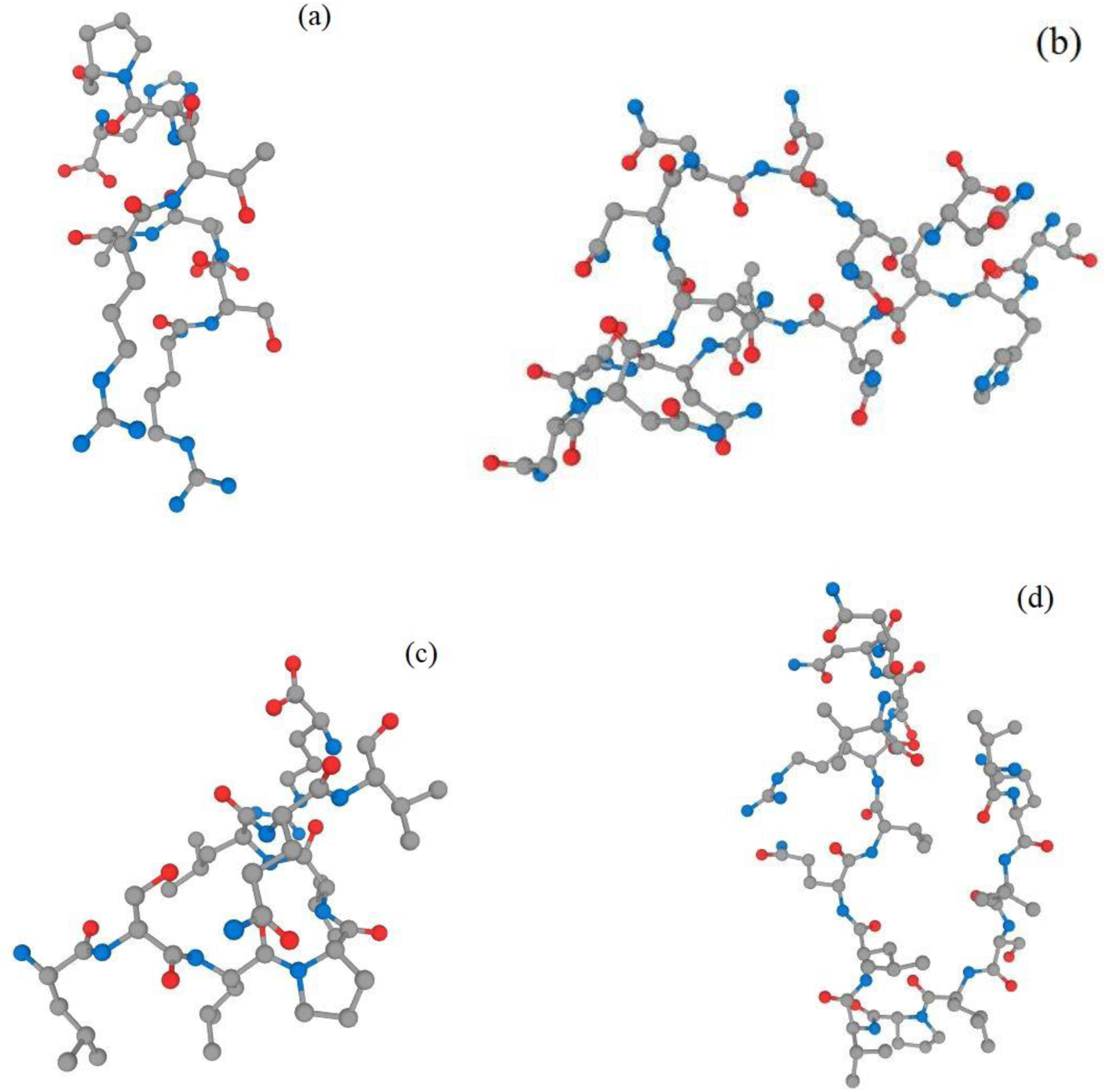
(a-d) 3D structure generation of the selected epitopes. (a) Selected T-cell MHC Class-I epitope for ORF1ab leader protein MT326102. (b) Selected T-cell MHC Class-II epitope for ORF1ab leader protein MT326102. (c) Selected T-cell MHC Class-I epitope for ORF1ab leader protein MT326175. (d) Selected T-cell MHC Class-II epitope for ORF1ab leader protein MT326175.

### Peptide-Protein Docking using HPEPDOCK Server

HPEPDOCK Server was used to perform docking of the peptide and protein. The purpose of the same was to analyse which of the two selected T-cell MHC Class-I epitope: **RSDARTAPH** and **LSLPVLQVR** had the lowest global energy. The epitope having the lowest global energy acts as a better vaccine candidate. The docking was performed against the HLA-A*11-01 allele whose *pdb* format was obtained through pre-docking using UCSF Chimera.

The global energy for the selected Class-I epitopes were: -182.706 and -191.198 respectively for **RSDARTAPH** and **LSLPVLQVR**. Out of the two MHC Class-I epitopes selected for leader proteins ORF1ab **MT326102** and **MT326175**, the global energy was lowest for **LSLPVLQVR**.

**Figure 4.**
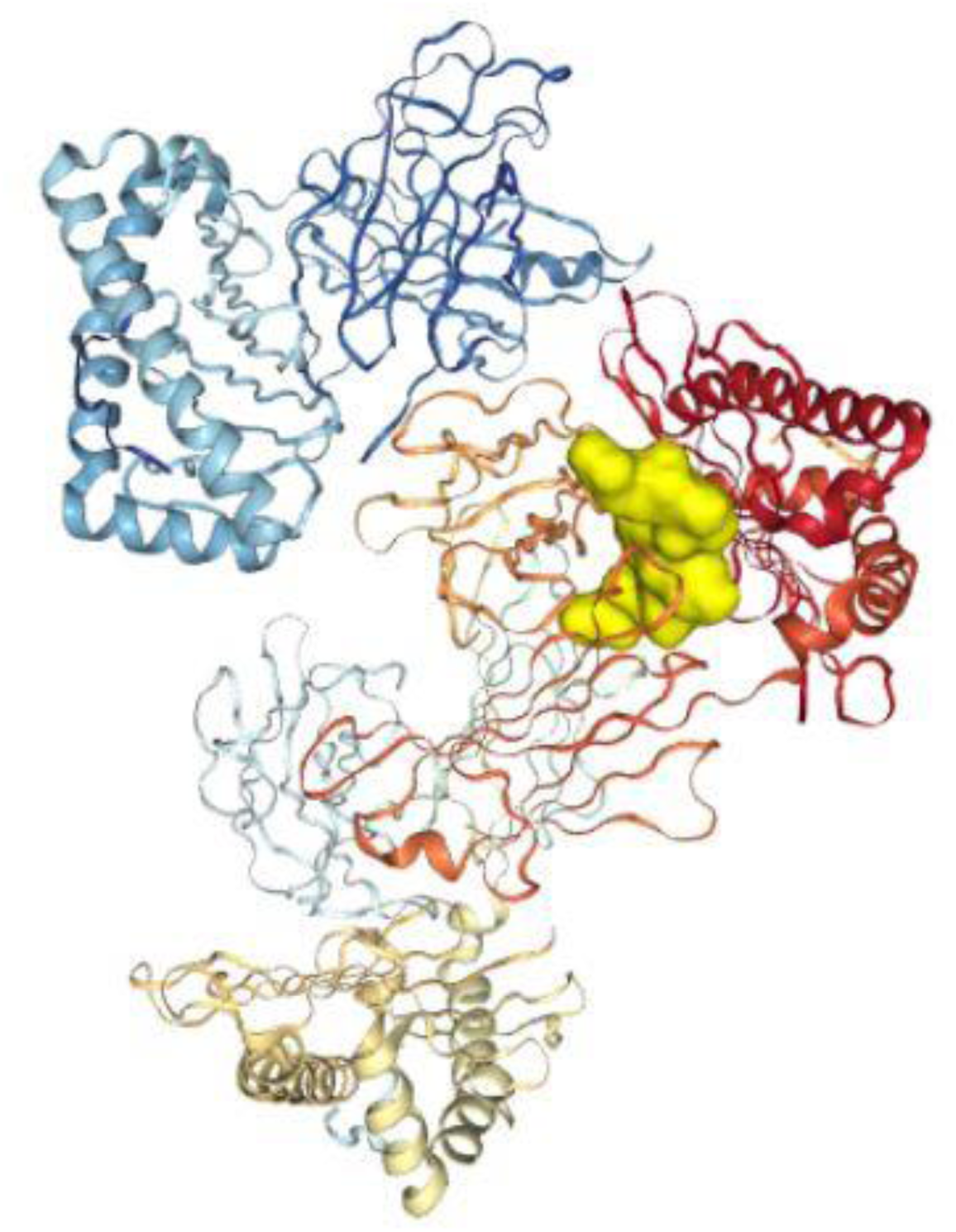
The peptide LSLPVLQVR is shown in the surface (yellow) format which has been docked against the HLA-A*11-01, displayed in the ribbon format. The format of the allele and the peptide are differed on purpose to show distinction.

For the MHC Class-II T-cell epitope, the docking was performed against the HLA DRB1*04-01 allele. Same procedure was followed as mentioned before. The two selected epitopes after the previous analysis were: **VQLNNNNNN** and **VQLSLPVLQ**. The global energy for the selected epitopes were -203.369 and -238.196 respectively.

Out of the two T-cell MHC Class-II epitopes selected for the ORF1ab leader proteins **MT326102** and **MT326175**, the lowest global energy was of **VQLSLPVLQ**.

**Figure 5.**
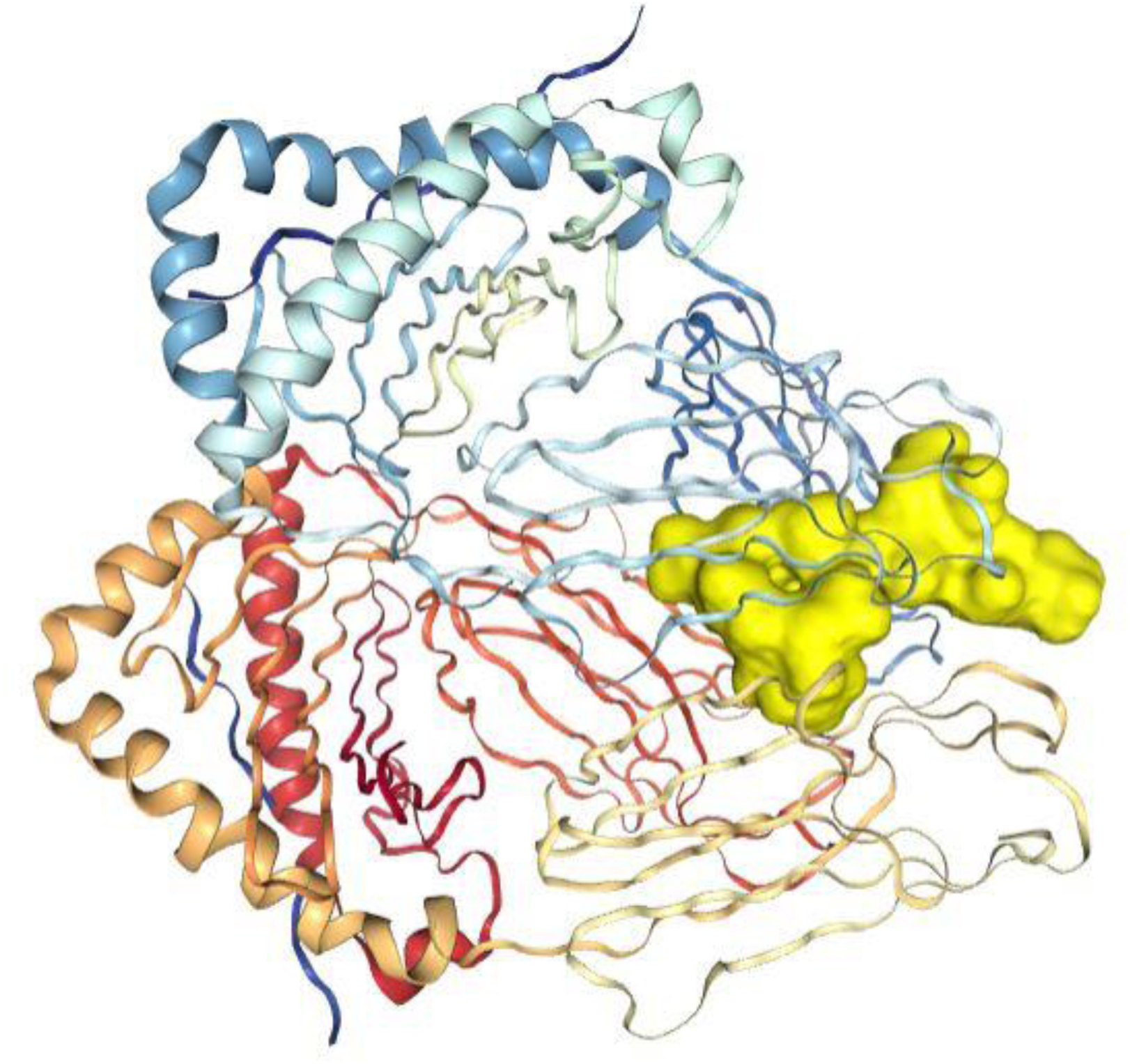
The peptide VQLSLPVLQ is shown in the surface (yellow) format which has been docked against the HLA DRB1*04-01, displayed in the ribbon format. The format of the allele and the peptide are differed on purpose to show distinction

## Results and Discussion

The scope of this study involved performing an *in-silica* analysis of the SARS-CoV-2 viral strain for the country of USA against human host. The ViPR database was used for the same to obtain all the protein sequences. A total of 14534 viral protein sequences were obtained whose extensive physicochemical analysis was done to select a group of two. This extensive analysis was performed using *Peptides* package in the R Software.

It was revealed that the two leader proteins ORF1ab **MT326102** and **MT326715** had the highest extinction coefficient and the lowest score on the GRAVY. Along with the given parameters, these leader proteins were highly stable and were also antigenic in nature.

The FASTA-formatted files of these selected proteins were taken and analysed to obtain the potential T-cell and B-cell epitopes. The T-cell epitopes of MHC Class-1 and MHC Class-II were analysed on the basis of their scores. Ten randomly selected T-cell epitopes from both the classes were taken for further analysis of allergenicity, toxicity, conservancy scores and antigenicity. Only one epitope from both the classes was selected which possessed a higher conservancy score (more than 90%), was non-toxic, non-allergic and antigenic in nature.

The two selected epitopes were then docked against their respective alleles to obtain the global energy scores. The epitopes which displayed the lowest global energy score on docking with the alleles were selected and proposed as successful and potential vaccine candidates

## Compliance with Ethical Standards

### Ethical Approval

The study doesn’t contain any work/experiment performed with human participants or animals done by the author.

